## Supplementary Information for "Intermittent precipitation and spatial Allee effects drive irregular vegetation patterns in semiarid ecosystems"

[1] *Department of Ecology and Complexity, Centre d'Estudis Avançats de Blanes (CEAB-CSIC), Blanes, Girona, Spain*
[2] *Instituto de Física Interdisciplinar y Sistemas Complejos (IFISC, CSIC-UIB), Campus UIB, 07122 Palma de Mallorca, Spain*
[3] *Department of Mathematics and Statistics, Utah State University, Logan, Utah*
[4] *Department of Mathematics, University of British Columbia, Vancouver, BC, Canada*
[*] *Corresponding author*


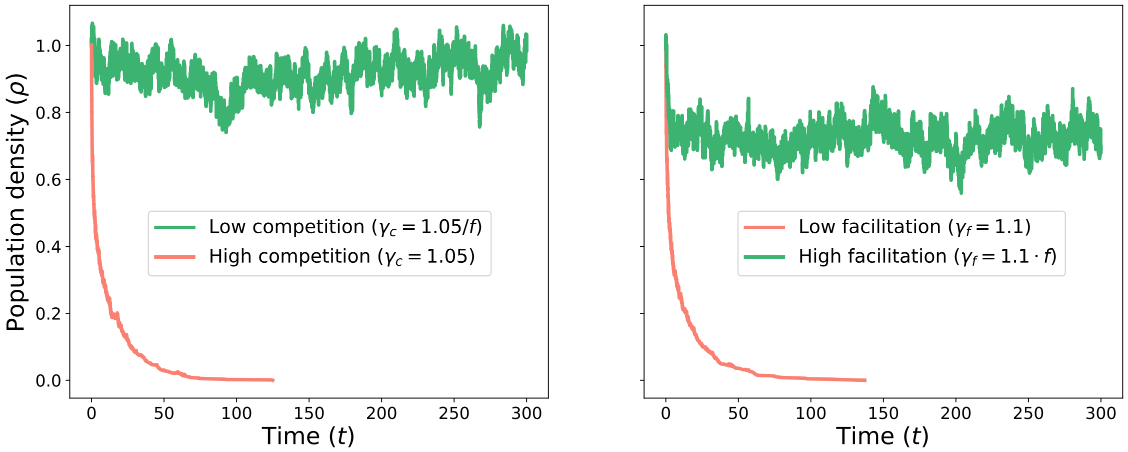


**Figure 1:** Example time series of a single realization of the Allee-based individual-based model in the absence of precipitation intermittency. Population density evolves under constant low (green) and high (red) competition or facilitation regimes. In the full stochastic model, precipitation events are implemented by switching between these two regimes through changes in the interaction kernel.

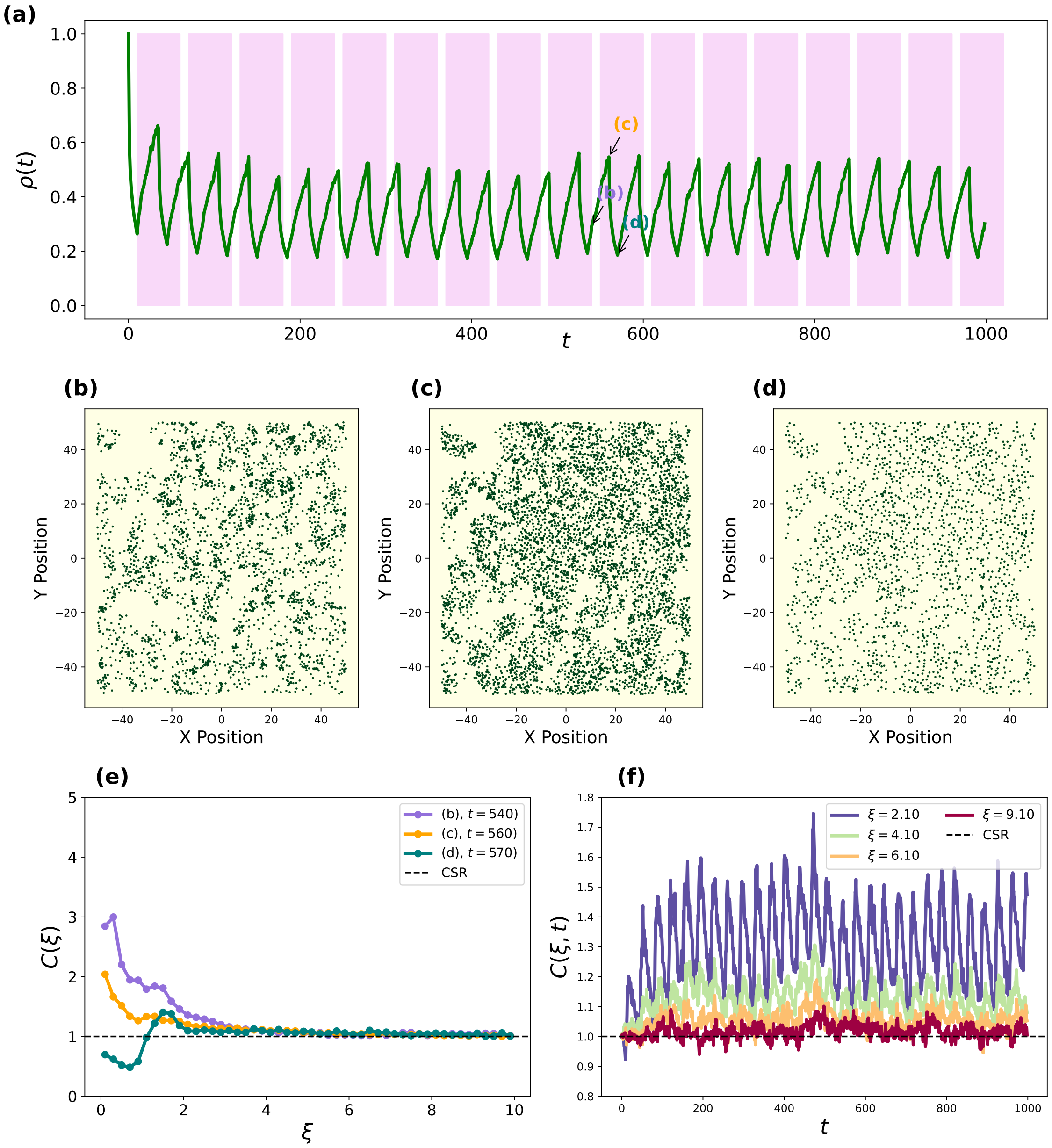


**Figure 2: Emergence and quantification of spatial clustering in a representative realization of the model with precipitation coupled to facilitation.** (a) Representative trajectory of population density as function of time with $d = 25$ and $\lambda = 0.1$. Pink regions correspond to time periods of enhanced facilitation due to precipitation events. (b)–(d) Snapshots of individual spatial configurations at successive times, showing the formation and reorganization of vegetation clusters. (e) Pair-correlation function computed at the times indicated in panels (b)–(d), compared with complete spatial randomness (CSR). (f) Time evolution of the pair-correlation function for different distances, illustrating the development and persistence of spatial structure relative to CSR.

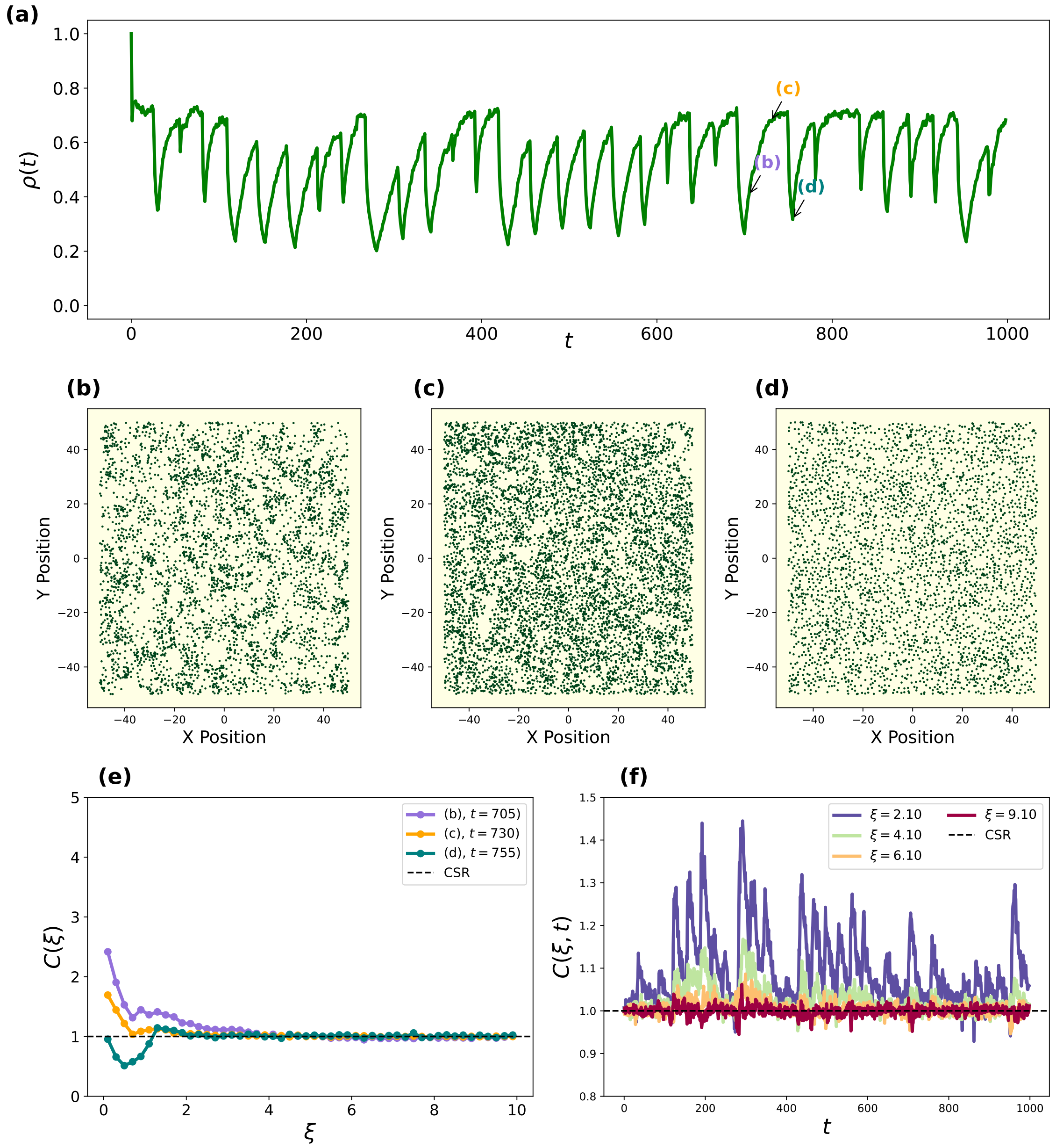


**Figure 3: Emergence and quantification of spatial clustering in a representative realization of the model with precipitation coupled to facilitation and stochastic precipitation events.** (a) Representative trajectory of population density as function of time with $d = 25$ and $\lambda = 0.2$. (b)–(d) Snapshots of individual spatial configurations at successive times, showing the formation and reorganization of vegetation clusters. (e) Pair-correlation function computed at the times indicated in panels (b)–(d), compared with complete spatial randomness (CSR). (f) Time evolution of the pair-correlation function for different distances, illustrating the development and persistence of spatial structure relative to CSR.

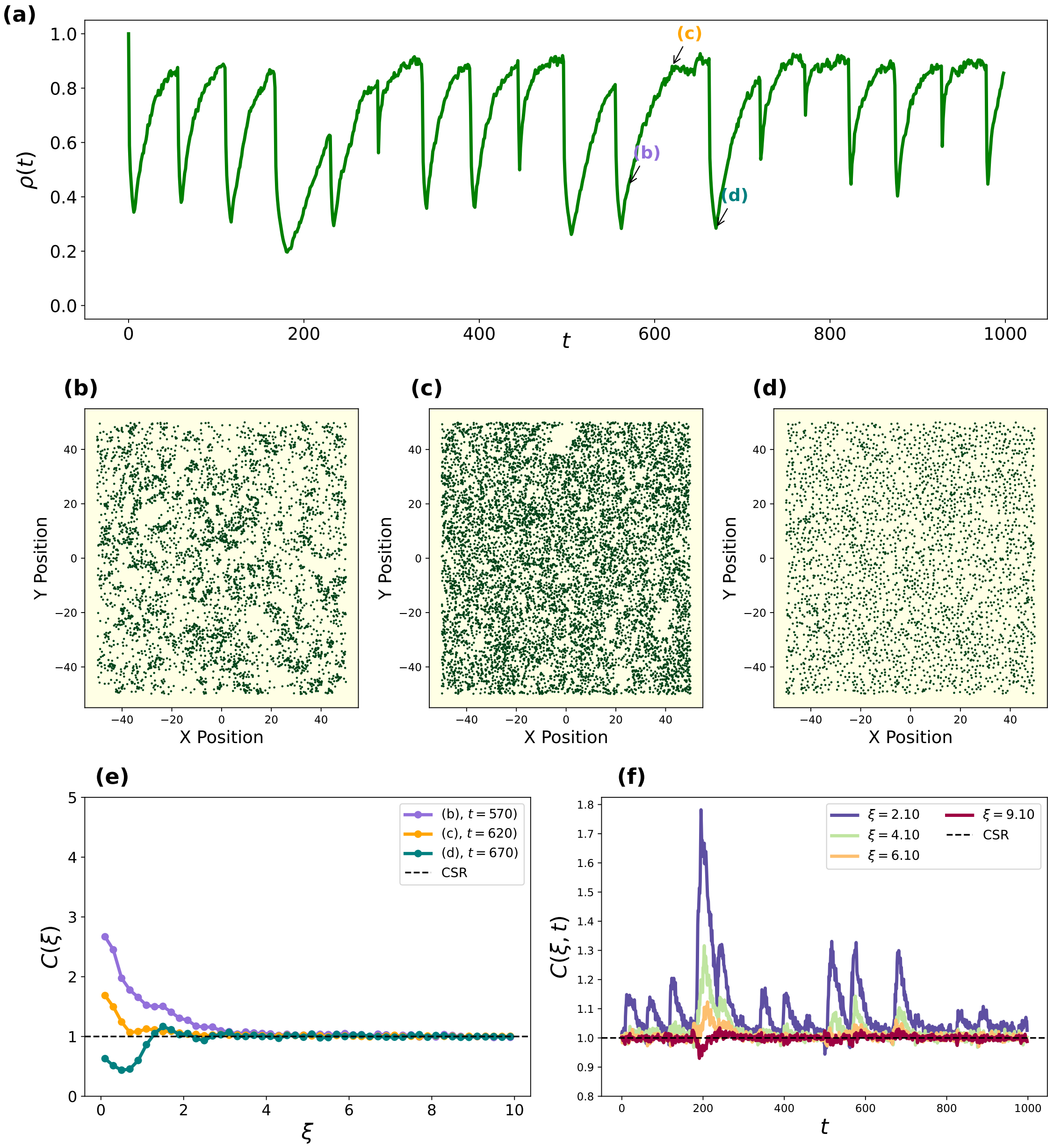


**Figure 4: Emergence and quantification of spatial clustering in a representative realization of the model with precipitation coupled to competition and stochastic precipitation events.** (a) Representative trajectory of population density as function of time with $d = 50$ and $\lambda = 0.2$. (b)–(d) Snapshots of individual spatial configurations at successive times, showing the formation and reorganization of vegetation clusters. (e) Pair-correlation function computed at the times indicated in panels (b)–(d), compared with complete spatial randomness (CSR). (f) Time evolution of the pair-correlation function for different distances, illustrating the development and persistence of spatial structure relative to CSR.

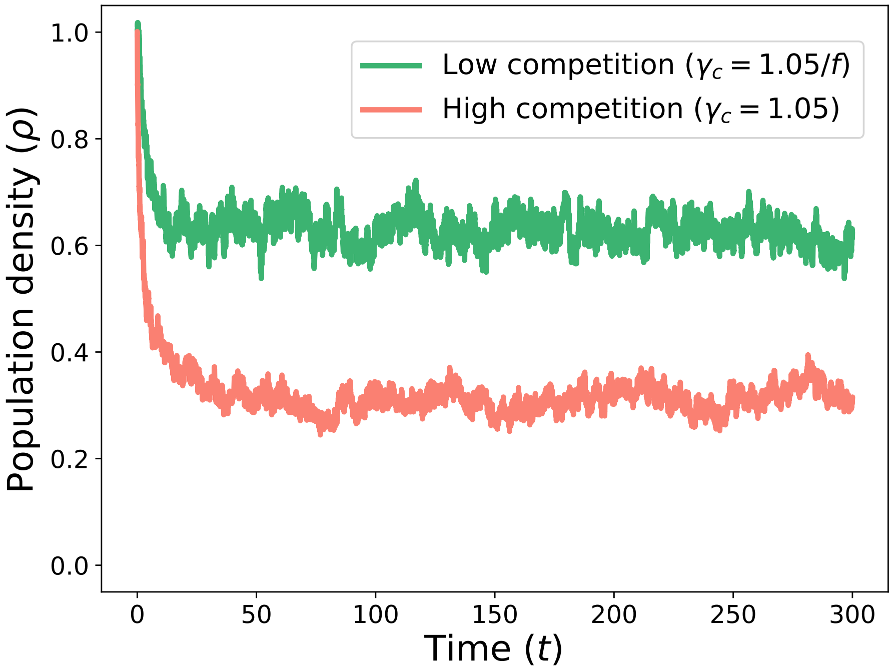


**Figure 5:** Example time series of a single realization of the logistic (non-Allee) model without precipitation intermittency, shown for constant low (green) and high (red) competition regimes. In simulations with precipitation events, the system alternates between these regimes by switching the competition kernel, analogously to the Allee-based model.

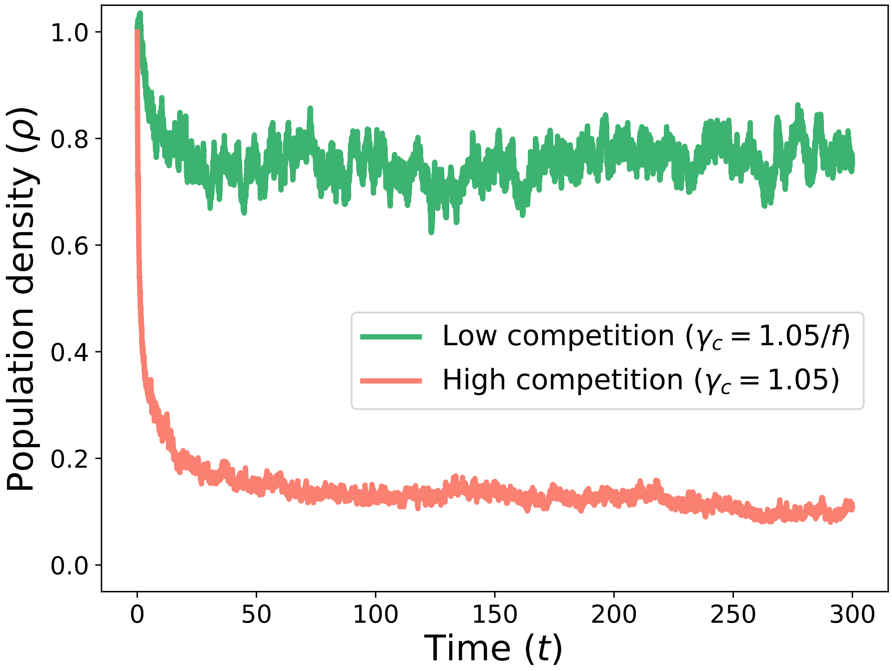


**Figure 6:** Example time series of a single realization of the logistic (non-Allee) model without precipitation intermittency, shown for constant low (green) and high (red) competition regimes with $\gamma_c = 1.8$ and $\gamma_{c_-} = 0.6$. In simulations with precipitation events, the system alternates between these regimes by switching the competition kernel, analogously to the Allee-based model.

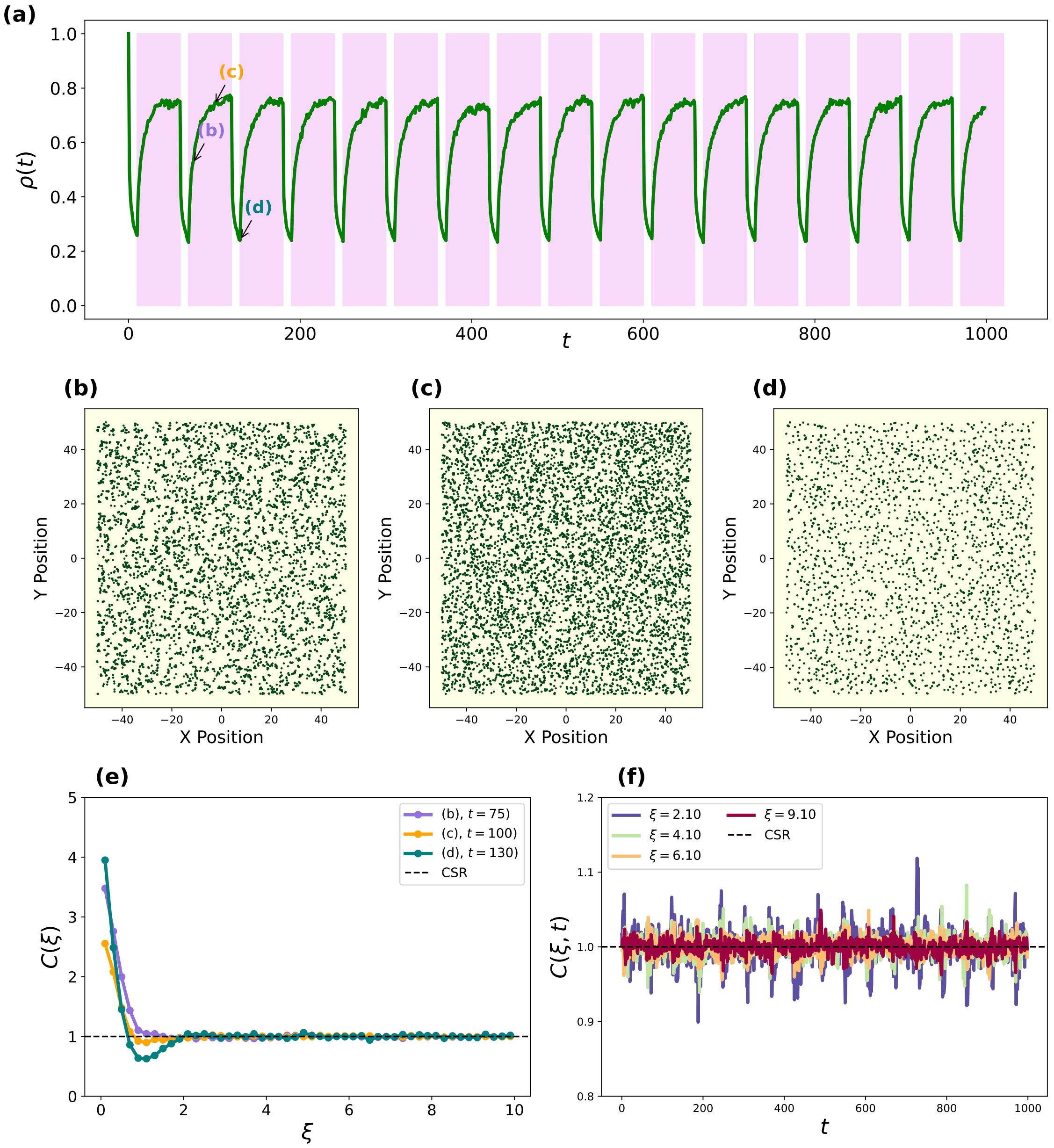


**Figure 7: Absence of large-scale clustering in the logistic model under intermittent precipitation with $\gamma_c = 1.8$ and $\gamma_{c_-} = 0.6$.** (a) Population density trajectory using a logistic model. (b–d) Spatial snapshots showing a near-uniform distribution of individuals; while small, transient groups appear due to short-range dispersal, no large-scale clusters emerge. (e) The PCF remains near 1 across most distances, suggesting little spatial correlation. (f) Time-resolved PCF confirms that without positive density-dependent feedback, vegetation cannot self-organize into structured patterns despite environmental fluctuations.

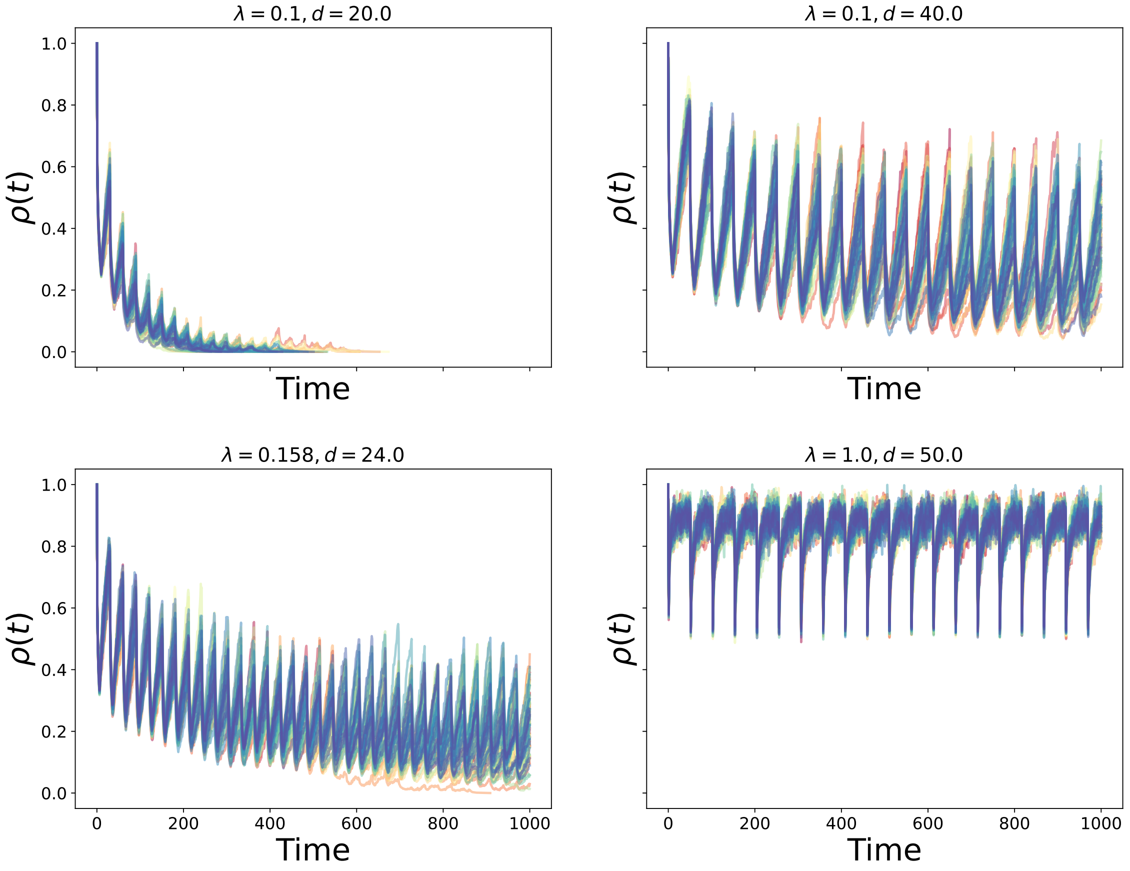


**Figure 8:** Individual realizations of the Allee-based model for different combinations of precipitation switching rate $\lambda$ and mean residence time $d$, with system size $L = 50$. Precipitation intermittency is coupled to competition. The panels illustrate how changes in the frequency and duration of favorable periods affect population persistence and temporal variability.

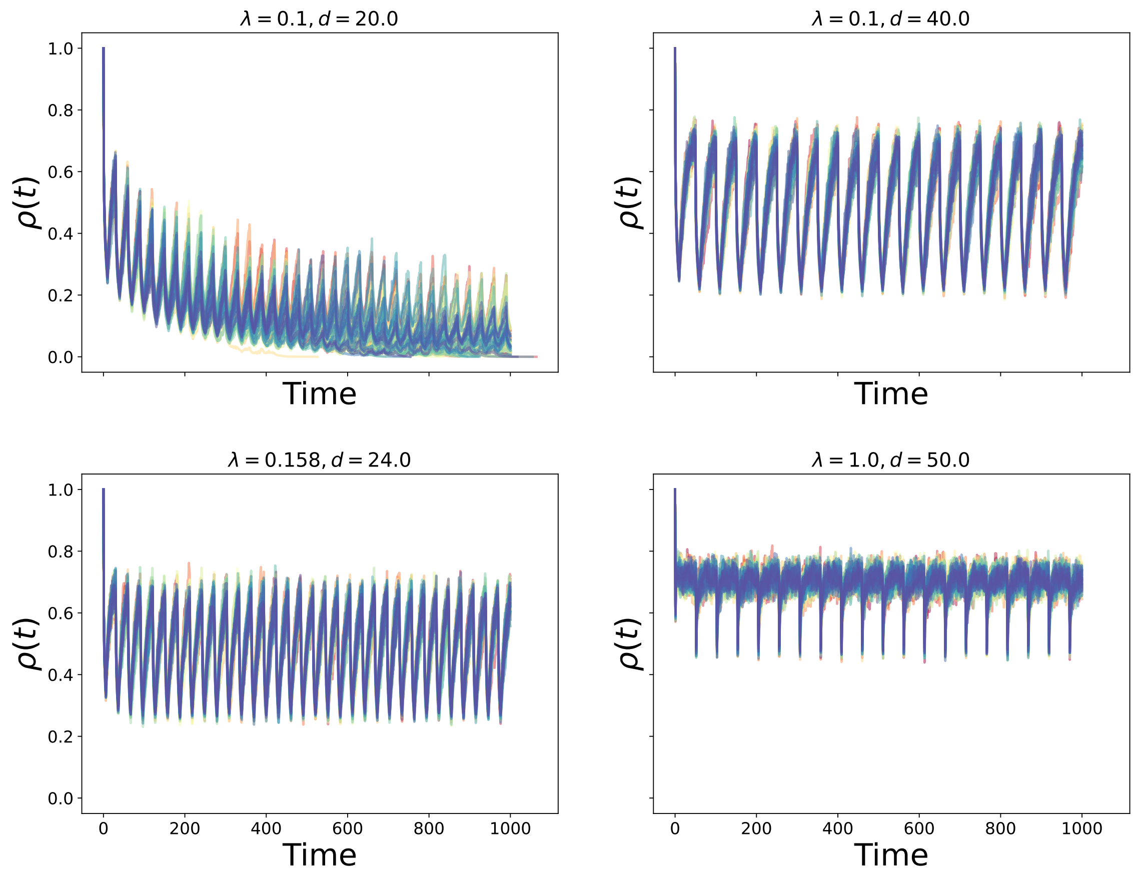


**Figure 9:** Individual realizations of the Allee-based model for the same parameter combinations as in Fig. 8, but with precipitation intermittency coupled to facilitation. Comparison with Fig. 8 highlights that the qualitative effects of intermittency on persistence and variability are similar across coupling mechanisms, despite differences in the underlying interaction pathways

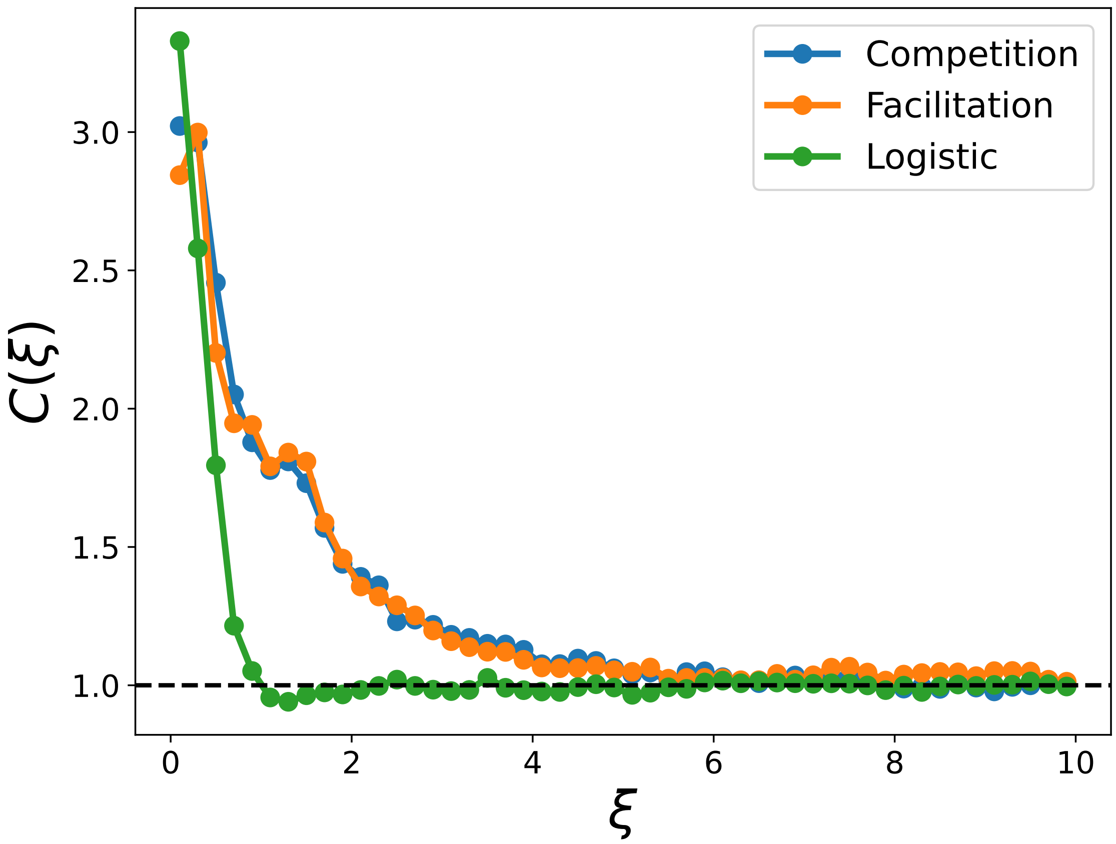


**Figure 10:** Pair-correlation functions (PCFs) obtained from the Allee model when precipitation modulates either competition (blue) or facilitation (orange), compared with the corresponding PCF of the logistic model (green). In the Allee model, the PCFs are nearly indistinguishable regardless of whether precipitation acts on facilitation or competition, indicating that spatial structure is robust to the specific coupling mechanism. By contrast, the logistic model exhibits a markedly different spatial signature, with a more rapid decay of correlations and a slight undershoot below unity, reflecting weaker clustering and the absence of positive density dependence.
